# Trace metal imaging of sulfate-reducing bacteria and methanogenic archaea at single-cell resolution by synchrotron X-ray fluorescence imaging

**DOI:** 10.1101/087585

**Authors:** Jennifer B. Glass, Si Chen, Katherine S. Dawson, Damian R. Horton, Stefan Vogt, Ellery D. Ingall, Benjamin S. Twining, Victoria J. Orphan

## Abstract

Metal cofactors are required for many enzymes in anaerobic microbial respiration. This study examined iron, cobalt, nickel, copper, and zinc in cellular and abiotic phases at the single-cell scale for a sulfate-reducing bacterium (*Desulfococcus multivorans*) and a methanogenic archaeon (*Methanosarcina acetivorans*) using synchrotron x-ray fluorescence microscopy. Relative abundances of cellular metals were also measured by inductively coupled plasma mass spectrometry. For both species, zinc and iron were consistently the most abundant cellular metals. *M. acetivorans* contained higher nickel and cobalt content than *D. multivorans*, likely due to elevated metal requirements for methylotrophic methanogenesis. Cocultures contained spheroid zinc sulfides and cobalt/copper-sulfides.

## Introduction

In anoxic natural and engineered environments, sulfate-reducing bacteria and methanogenic archaea perform the last two steps of organic carbon respiration, releasing sulfide and methane. Sulfate-reducing bacteria and methanogenic archaea can exhibit cooperative or competitive interactions depending on sulfate and electron donor availability (Brileya et al. 2014; Bryant et al. 1977; Ozuolmez et al. 2015; Stams and Plugge 2009). Methanol (CH_3_OH), the simplest alcohol, is an important substrate for industrial applications (Bertau et al.2014) and microbial metabolisms. In the presence of methanol, sulfate reduction and methanogenesis occur simultaneously in cocultures (Dawson et al. 2015; Phelps et al. 1985), anoxic sediments (Finke et al. 2007; Oremland and Polcin 1982), and anaerobic digesters (Spanjers et al. 2002; Weijma and Stams 2001). Methanol has also been studied as a substrate for stimulating organochlorine degradation in sediment reactors containing sulfate-reducing bacteria and methanogenic archaea (Drzyzga et al. 2002).

Metalloenzymes are essential for both sulfate reduction and methylotrophic methanogenesis (Barton et al. 2007; Ferry 2010; Glass and Orphan 2012; Thauer et al. 2010). Iron is needed for cytochromes and iron-sulfur proteins in both types of organisms (Fauque and Barton 2012; Pereira et al. 2011; Thauer et al. 2008). Cobalt and zinc are present in the first enzymes in sulfate reduction (ATP sulfurylase, Sat; Gavel et al. 1998; Gavel et al. 2008), and methylotrophic methanogenesis (methanol:coenzyme M methyltransferase; Hagemeier et al. 2006). Nickel is found in the final enzyme in methanogenesis (methyl coenzyme M reductase; Ermler et al. 1997), and zinc is present in the heterodisulfide reductase that recycles cofactors for the methyl coenzyme M reductase enzyme (Hamann et al. 2007). Nickel and cobalt are required by methanogenic archaea and sulfate-reducing bacteria that are capable of complete organic carbon oxidization for carbon monoxide dehydrogenase/acetyl Co-A synthase in the Wood-Ljungdahl CO_2_ fixation pathway (Berg 2011; Ragsdale and Kumar 1996). Hydrogenases containing Ni and Fe are functional in many, but not all, sulfate-reducing bacteria (Osburn et al. 2016; Pereira et al. 2011) and methylotrophic methanogens (Guss et al. 2009; Thauer et al. 2010). Evidence for high metabolic metal demands is provided by limited growth of methanogenic archaea without Co and Ni supplementation in methanol-fed monocultures (Scherer and Sahm 1981) and anaerobic bioreactors (Florencio et al. 1994; Gonzalez-Gil et al. 1999; Paulo et al. 2004; Zandvoort et al. 2003; Zandvoort et al. 2006).

Sulfate-reducing bacteria produce sulfide, which can remove toxic metals from contaminated ecosystems due to precipitation of metal sulfides with low solubility (Paulo et al 2015). Metal sulfides may also limit the availability of essential trace metals for microbial metabolism (Glass and Orphan 2012; Glass et al. 2014). In sulfidic environments such as marine sediments and anaerobic digesters, dissolved Co and Ni are present in nanomolar concentrations (Glass et al. 2014; Jansen et al. 2005). These metals are predominantly present as solid metal sulfide precipitates (Drzyzga et al. 2002; Luther III and Rickard 2005; Moreau et al. 2013) and/or sorbed to anaerobic sludge (van Hullebusch et al. 2006; van Hullebusch et al. 2005; van Hullebusch et al. 2004). The bioavailability of metals in these solid phases to anaerobic microbes remains relatively unknown. Previous studies suggest that methanogenic archaea can leach Ni from silicate minerals (Hausrath et al. 2007) and metal sulfides (Gonzalez-Gil et al. 1999; Jansen et al. 2007). Sulfidic/methanogenic bioreactors (Jansen et al. 2005) and *D. multivorans* monocultures (Bridge et al. 1999) contain high-affinity Co-/Ni- and Cu-/Zn-binding ligands, respectively, which may aid in liberating metal micronutrients from solid phases when they become growth-limiting.

Due to the importance of trace metals for anaerobic microbial metabolisms in bioremediation and wastewater treatment, extensive efforts have focused on optimizing metal concentrations to promote microbial organic degradation in anaerobic digesters (for review, see Demirel and Scherer (2011)). Numerous studies have investigated the effect of heavy metals on anaerobic metabolisms at millimolar concentrations in heavy-metal contaminated industrial wastewaters, whereas few studies have investigated interactions between anaerobic microbes and transition metals at the low micro- to nanomolar metal concentrations present in most natural ecosystems and municipal wastewaters (see Paulo et al. 2015 for review). Studies of the metal content of anaerobic microbes have primarily measured monocultures using non-spatially resolved techniques such as ICP-MS (Barton et al. 2007; Cvetkovic et al. 2010;Scherer et al. 1983). Little is known about the effect of coculturing on cellular elemental composition and mineralogy due to changes in geochemistry (e.g. via sulfide production) of the medium and/or microbial metabolisms (e.g. via competition for growth-limiting substrates).

In this study, we measured cellular elemental contents and imaged extracellular metallic minerals for sulfate-reducing bacteria and methanogenic archaea grown in mono- and co-culture. For the model sulfate-reducing bacterium, we chose the metabolically versatile species *Desulfococcus multivorans*, which is capable of complete organic carbon oxidation. *Methanosarcina acetivorans* C2A, a well-studied strain capable of growing via aceticlastic and methylotrophic methanogenesis, but not on H_2_/CO_2_, was selected as the model methanogenic archaeon. These species were chosen because they are the most phylogenetically similar to pure culture isolates available to syntrophic consortia of anaerobic methanotrophic euryarchaeota (ANME-2) and sulfate-reducing bacteria (Desulfosarcina/Desulfococcus) partner that catalyze the anaerobic oxidation of methane in marine sediments (see Dawson et al. (2015) for more on coculture design). Individual cells of mono- and cocultures of these two species were imaged for elemental content on the Bionanoprobe (Chen et al. 2013) at the Advanced Photon Source (Argonne National Laboratory) and measured for relative abundance of bulk cellular metals by ICP-MS.

## Materials and Methods

### Culture growth conditions

The growth medium contained (in g L^−1^): NaCl, 23.4; MgSO_4_ 7H_2_O, 9.44; NaHCO_3_, 5.0; KCl, 0.8; NH_4_Cl, 1.0; Na_2_HPO_4_, 0.6; CaCl_2_ 2H_2_O, 0.14; cysteine-HCl, 0.25; resazurin, 0.001, 3 × 10^−6^ Na_2_ SeO_3_ supplemented with DSM-141 vitamin (including 1 μg L^−1^ vitamin B_12_) and trace element solutions containing metal concentrations (provided below as measured by ICP-MS) and 1.5 mg L^−1^ nitrilotriacetic acid (Atlas 2010). The medium (pH 7.6) was filter sterilized in an anoxic chamber (97% N_2_ and 3% H_2_ headspace) and reduced with 1 mM Na_2_S.

*Monocultures of Desulfococcus multivorans* (DSM 2059) and Methanosarcina acetivorans strain C2A (DSM 2834) were inoculated into 20 mL culture tubes containing 10 mL of media with N_2_:CO_2_ (80:20) headspace, and sealed with butyl rubber stoppers and aluminum crimp seals. *D. multivorans* monocultures were amended with filter-sterilized lactate (20 mM). *M. acetivorans* monocultures were amended with filter-sterilized methanol (66 mM). Equal proportions of dense monocultures in early stationary stage (as assessed by OD600 measurements; Fig. S1) were inoculated into sterile media and amended with filter-sterilized lactate (20 mM) and methanol (66 mM) to form the coculture. Cultures were grown at 30°C without shaking. After 12 days of growth (Fig. S1), mono- and cocultures were pelleted and frozen for ICP-MS analysis, or prepared for SXRF imaging.

### Fluorescence in situ hybridization

In order to confirm that monocultures were free of contamination, and to determine the relative abundance of *D. multivorans* and *M. acetivorans* in coculture, fluorescence in *situ* hybridization (FISH) was performed on separate aliquots from the same time point of the cell culture used for SXRF analyses. One mL of cell culture was preserved in 3% paraformaldehyde for 1-3 hours, then washed and resuspended in 200 μΤ of 3x PBS:ethanol as described in Dawson et al. (2012). Four microliters of fixed cells were spotted onto a glass slide and hybridized with an oligonucleotide probe targeting *Methanosarcina acetivorans* MSMX860 (Raskin et al. 1994) and the deltaproteobacterial probe Delta495a (Loy et al. 2002) and cDelta495a (Macalady et al. 2006). The FISH hybridization buffer contained 45% formamide, and the hybridization was carried out at 46°C for 2 hours followed by a 15 minute wash in 48°C washing buffer (Daims et al. 2005). The slides were rinsed briefly in distilled water, and mounted in a solution of DAPI (5 μg/mL) in Citifluor AF-1 (Electron Microscopy Services). Imaging was performed with a 100x oil immersion objective (Olympus PlanApo). Cell counts were performed by hand. Multiple fields of view from replicate wells were compiled and counted on the basis of fluorescence in DAPI (all cells), Cy3 (bacteria), and FITC (archaea).

### ICP-MS

Frozen cell pellets were dried (yielding ∼4 mg dry weight per sample) in acid-washed Savillex Teflon vials in a laminar flow hood connected to ductwork for exhausting acid fumes. Cells were digested overnight at 150°C in 2 mL of trace metal grade nitric acid and 200 μL hydrogen peroxide, dried again, and dissolved in 5 mL 5% nitric acid. The medium was diluted 1:50 in nitric acid. The elemental content of microbial cells and media was analyzed by ICP-MS (Element-2, University of Maine Climate Change Institute). Sterile medium contained the following concentrations (in μM): P, 800; Zn, 7; Fe, 4; Co, 2; Ni, 0.9; Cu, 0.3. Digestion acid blanks contained (in nM): P, 127; Zn, 12; Fe, 5; Co, 0.007; Ni, 0.9; Mo, 0.02; Cu, 0.1; V, 0.03.

### SXRF sample preparation

Monocultures were prepared for SXRF analysis without chemical fixation by spotting onto silicon nitride (SiN) wafers (Silson Ltd., cat. 11107126) followed by rinsing with 10 mM HEPES buffer (pH 7.8). To enable FISH microscopy after SXRF analysis, cocultures were chemically preserved prior to analysis by incubation on ice for 1 hour in 50 mM HEPES and 0.6 M NaCl (pH 7.2) containing 3.8% paraformaldehyde and 0.1% glutaraldehyde that had been cleaned of potential trace-metal contaminants with cation exchange resin (Dowex 50-W X8) using established protocols (Price et al. 1988; Twining et al. 2003). Cells were then centrifuged, re-suspended in 10 mM HEPES buffer (pH 7.8) and either embedded in resin and thin sectioned following the methods described in McGlynn et al. (2015) or spotted directly onto SiN wafers.

### SXRF analyses

Whereas ICP-MS measurements cannot delineate the elemental contributions of co-occurring cell types, SXRF imaging enables elemental quantification of the specific cell of interest (Fahrni 2007; Ingall et al. 2013; Kemner et al. 2004; Nuester et al. 2012; Twining et al. 2003; Twining et al. 2008). SXRF analyses were performed at the Bionanoprobe (beamline 21-ID-D, Advanced Photon Source, Argonne National Laboratory). Silicon nitride wafers were mounted perpendicular to the beam as described in Chen et al. (2013). SXRF mapping was performed with monochromatic 10 keV hard X-rays focused to a spot size of ∼100 nm using Fresnel zone plates. Concentrations and distributions of all elements from P to Zn were analyzed in fine scans using a step size of 100 nm and a dwell time of 150 ms. An X-ray fluorescence thin film (AXO DRESDEN, RF8-200-S2453) was measured with the same beamline setup as a reference. MAPS software was used for per-pixel spectrum fitting and elemental content quantification (Vogt 2003). Sample elemental contents were computed by comparing fluorescence measurements with a calibration curve derived from measurements of a reference thin film.

Regions of interest (ROIs) were selected with MAPS software by highlighting each microbial cell (identified based on elevated P content with care taken to avoid regions of elevated non-cellular metals) or particle (identified based on elevated metal content). Each ROI (n=14 and n=17 for *D. multivorans* (radius: 0.60 ± 0.01 μm) and *M. acetivorans* (radius: 0.48 ± 0.01 μm), respectively, and n=13 for the coculture (radius: 0.96 ± 0.01 μm)) was background corrected to remove elements originating from each section of the SiN grid on which cells were spotted. To do so, the mean of triplicate measurements of area-normalized elemental content for blank areas bordering the analyzed cells was subtracted from cellular ROIs. The background-corrected area-normalized molar elemental content was then multiplied by cellular ROI area to obtain molar elemental content per cell, which was then divided by the cell volume (4/3πr^3^, assuming spherical cells) to yield total metal content per cell volume, in units of mmol L-1. Visualization of elemental co-localization was performed with MAPS software. Statistical analysis was performed with JMP Pro (v. 12.1.0) using the Tukey-Kramer HSD test.

**Table 1.** Mean and standard error (in parentheses) of elemental contents normalized per cellular volume and per cell as measured by SXRF. Monocultures were prepared without chemical fixation, and cocultures were prepared with paraformaldehyde and glutaraldehyde fixation, followed by spotting onto silicon nitride wafers as described in the text. A, B and C superscripts indicate statistically different element contents (p < 0.05 based on Tukey-Kramer HSD test).

| Culture | Substrate (mM) | P | S | Fe | Co | Ni | Cu | Zn |
| --- | --- | --- | --- | --- | --- | --- | --- | --- |
|  |  | Element per cellular volume (mmol L<sup>-1</sup>) |  |  |  |  |  |  |
| 100% *Methanosarcina acetivorans* DSM 2834 (n = 14) | Methanol (66 mM) | 382<sup>A</sup><br>(47) | 4553<sup>A</sup><br>(458) | 38<sup>A</sup><br>(4) | 3.1<sup>A</sup><br>(0.3) | 0.44<sup>A</sup><br>(0.04) | 11<sup>A</sup><br>(1) | 38<sup>A</sup><br>(3) |
| 100% *Desulfococcus multivorans* DSM 2059 (n = 17) | Lactate (20 mM) | 55<sup>B</sup><br>(7) | 96<sup>B</sup><br>(11) | 22<sup>B</sup><br>(4) | 0.5<sup>B</sup><br>(0.1) | 0.11<sup>B</sup><br>(0.02) | 3<sup>B</sup><br>(1) | 36<sup>A</sup><br>(10) |
| 77% *Methanosarcina acetivorans* DSM 2834, 23% *Desulfococcus multivorans* DSM 2059 (n = 13) | Methanol (66 mM), lactate (20 mM) | 353<sup>A</sup><br>(32) | 234<sup>B</sup><br>(19) | 3.6<sup>C</sup><br>(0.3) | 2.0<sup>C</sup><br>(0.2) | 0.15<sup>B</sup><br>(0.01) | 0.13<sup>C</sup><br>(0.04) | 2.4<sup>B</sup><br>(0.2) |
|  |  | Element per cell (mol x 10<sup>-18</sup> cell<sup>-1</sup>) |  |  |  |  |  |  |
| 100% *Methanosarcina acetivorans* DSM 2834 (n = 14) | Methanol (66 mM) | 178<sup>A</sup><br>(28) | 2167<sup>A</sup><br>(318) | 19<sup>A</sup><br>(4) | 1.5<sup>A</sup><br>(0.2) | 0.20<sup>A</sup><br>(0.03) | 5<sup>A</sup><br>(1) | 17<sup>AB</sup><br>(2) |
| 100% *Desulfococcus multivorans* DSM 2059 (n = 17) | Lactate (20 mM) | 60<sup>A</sup><br>(11) | 107<sup>B</sup><br>(19) | 24<sup>A</sup><br>(6) | 0.5<sup>A</sup><br>(0.1) | 0.12<sup>A</sup><br>(0.02) | 3<sup>A</sup><br>(1) | 39<sup>A</sup><br>(12) |
| 77% *Methanosarcina acetivorans* DSM 2834, 23% *Desulfococcus multivorans* DSM 2059 (n = 13) | Methanol (66 mM), lactate (20 mM) | 1252<sup>B</sup><br>(69) | 855<sup>C</sup><br>(69) | 13<sup>A</sup><br>(1) | 7<sup>B</sup><br>(1) | 2<sup>B</sup><br>(1) | 0.5<sup>B</sup><br>(0.2) | 9<sup>B</sup><br>(1) |

## Results

### Cellular elemental content of monocultures

Cellular metal contents of *M. acetivorans* and *D. multivorans* monocultures followed the trend Zn? Fe > Cu > Co > Ni when measured by SXRF, and Zn? Fe > Co > Ni > Cu when measured by ICP-MS (Fig. 1). When normalized to cell volume, cellular S measured by SXRF was 50x higher in methanol-grown *M. acetivorans* (n=14) than lactate-grown *D. multivorans* (n=17). Cellular P, Fe, Co, Ni and Cu were 4-7x higher in *M. acetivorans* than *D. multivorans*, and cellular Zn was not significantly different between the two microbes (Table 1).

**Figure 1.**
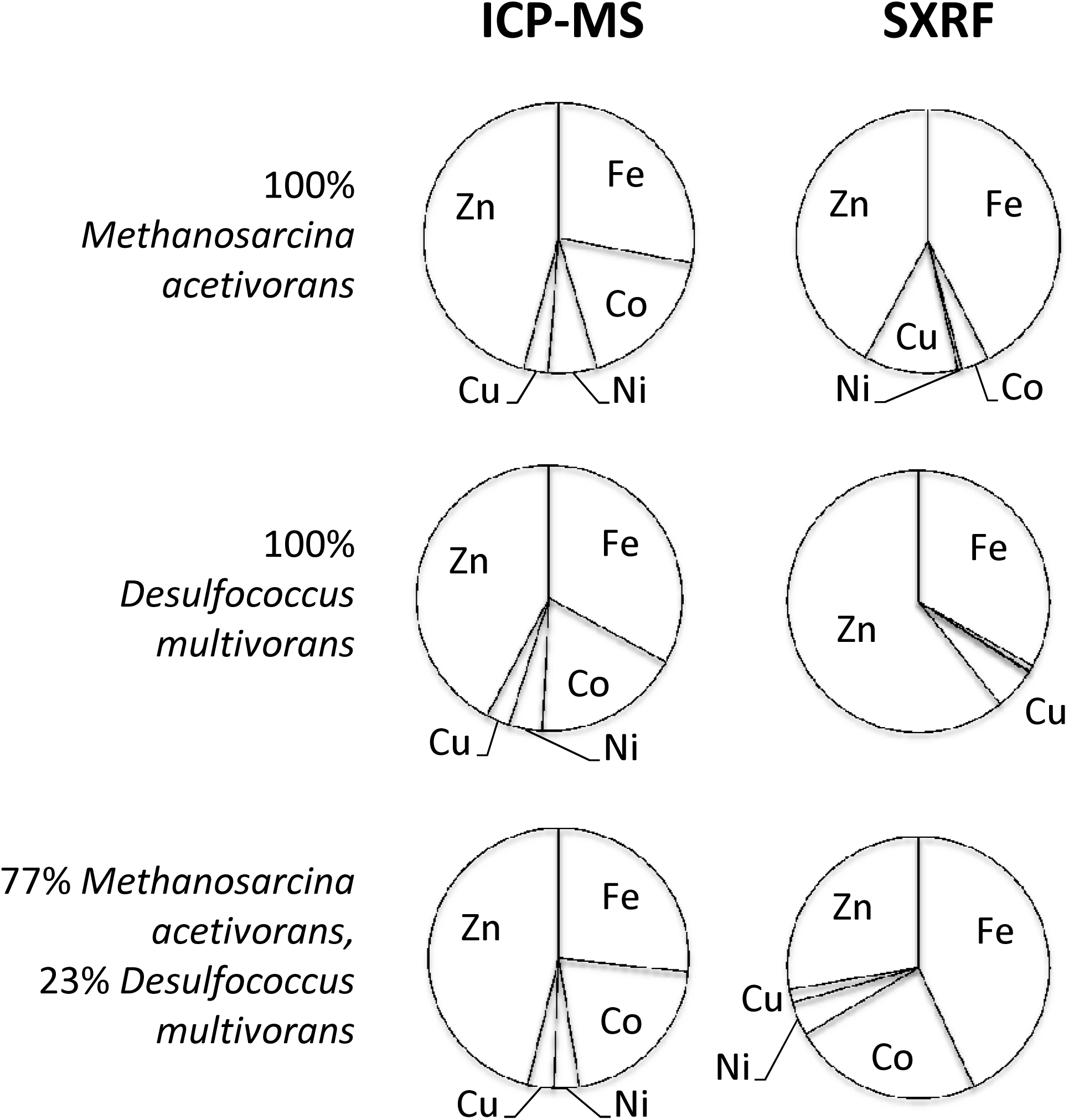
Proportions of each cellular metal (Fe, Co, Ni, Cu and Zn) for monocultures of *Methanosarcina acetivorans* (n=14), monocultures of *Desulfococcus multivorans* (n=18), and cocultures of 77% *M. acetivorans* and 23% *D. multivorans* (n=12) measured by ICP-MS (bulk measurement) and SXRF (single cell average).

### Relative abundance of species in coculture

Coculturing of both species for 12 days in media containing methanol and lactate resulted in dominance of *M. acetivorans* (77%, or 1,753 cells hybridized with the MSMX860 FISH probe) over *D. multivorans* (23%, or 522 cells hybridized with the Delta495a FISH probe) for 2,275 total cells counted in ten 100x (125 × 125 μm) fields of view. Cells were ∼1 μm^2^ cocci. No other cells exhibited DAPI staining other than those that hybridized with MSMX860 and Delta495a oligonucleotide probes. Attempts at FISH microscopy after SXRF analysis were unsuccessful due to x-ray radiation damage of the cells.

### Cellular elemental content of cocultures

ICP-MS measurements showed that the relative abundance of cellular metals remained relatively constant between mono- and cocultures, whereas SXRF data indicated that the coculture contained a relatively higher proportion of Co than the monocultures (Fig. 1). SXRF imaging showed no visual difference in elemental distribution between cells in the coculture (Fig. 2), although the relatively small size of the cells relative to the focused x-ray spot may have limited our ability to discern subtle differences. Cocultures, which were fixed with paraformaldehyde and glutaraldehyde for subsequent fluorescence microscopy, were larger (radius: 0.96 ± 0.01 μm) than monocultures (D. *multivorans* radius: 0.60 ± 0.01 μm; *M. acetivorans* radius: 0.48 ± 0.01 μm), which were not fixed prior to analysis. When normalized on a per cell basis, cocultures contained 5-20x higher P, Co and Ni than monocultures; however, when normalized to cellular volume, the larger cell volumes of the cocultures resulted in significantly less Fe, Cu and Zn per cellular volume than either of the monocultures (Table 1).

**Figure 2.**
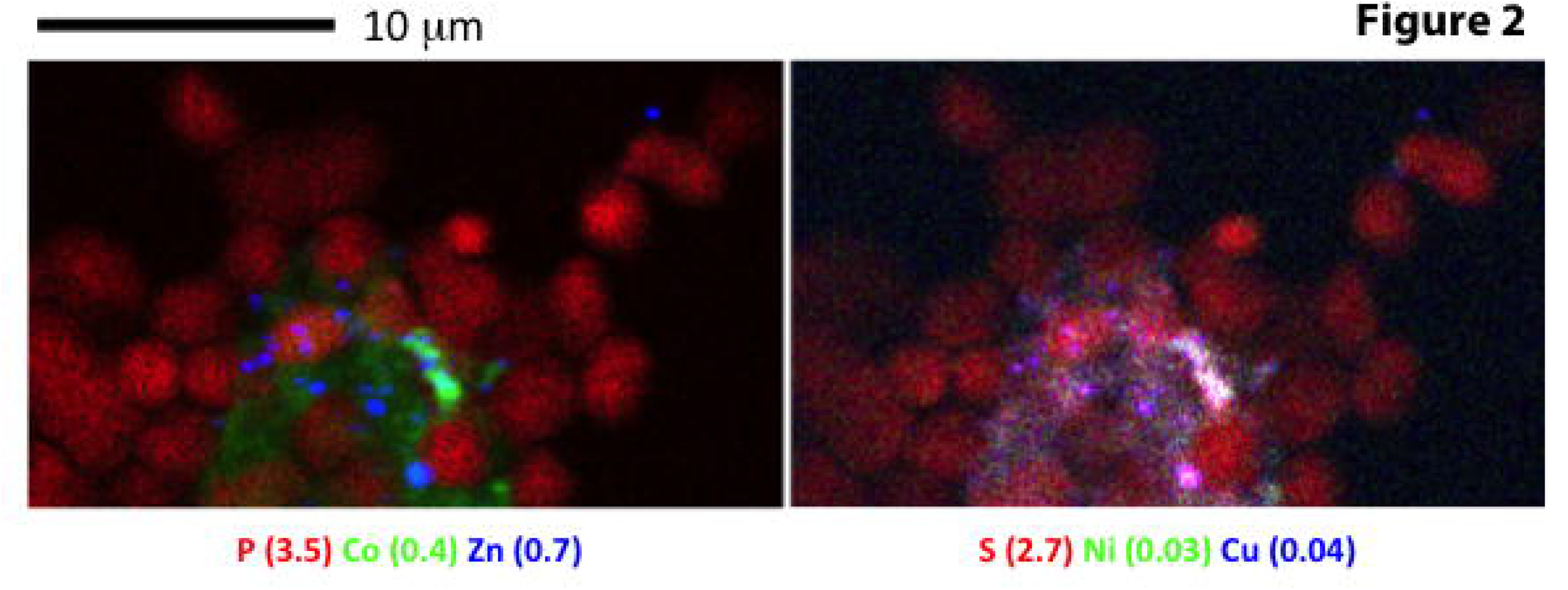
. SXRF co-localization of P (red), Co (green), and Zn (blue; left panel), and S (red), Ni (green), and Cu (blue; right panel) for whole cells of 77% *Methanosarcina acetivorans* and 23% *Desulfococcus multivorans* in coculture. Values in parentheses are maxima in μg cm^−2^ for each element.

### Non-cellular metals in cocultures

In whole cell SXRF images, ∼30 “hot spots” (discrete semi-circular areas with low-P and elevated metals, indicative of nano-sized minerals) of Zn (max: 0.7 μg cm^−2^), Co (max: 0.4 μg cm^−2^) and S (max: 2.7 μg cm^−2^) were present in the center of a cluster of ∼30 cocultured cells identified as P-containing cocci (Fig.2). In thin sections, semi-circular non-cellular small Zn hotspots (0.6 ± 0.1 μm^2^) containing ∼1:1 molar ratios of Zn:S (17 ± 2 μg Zn cm^−2^: 7.6 ± 0.7 μg S cm^−2^) were interspersed amongst cell clusters (n=8; Fig.3a-e) along with more numerous spheroid non-cellular Co hot spots of the same size (0.6 ± 0.1 μm^2^) containing 2.1 ± 0.1 μg Co cm^−2^, 3.4 ± 0.2 μg S cm^−2^, and 1.3 ± 0.1 μg Cu cm^−2^ (n=45; Fig.3a-e). Discrete semi-circular hot spots of elevated Ni (max: 2.9 μg cm^−2^) with low S were observed in two imaging fields (n=8; Fig.3b,c).

**Figure 3.**
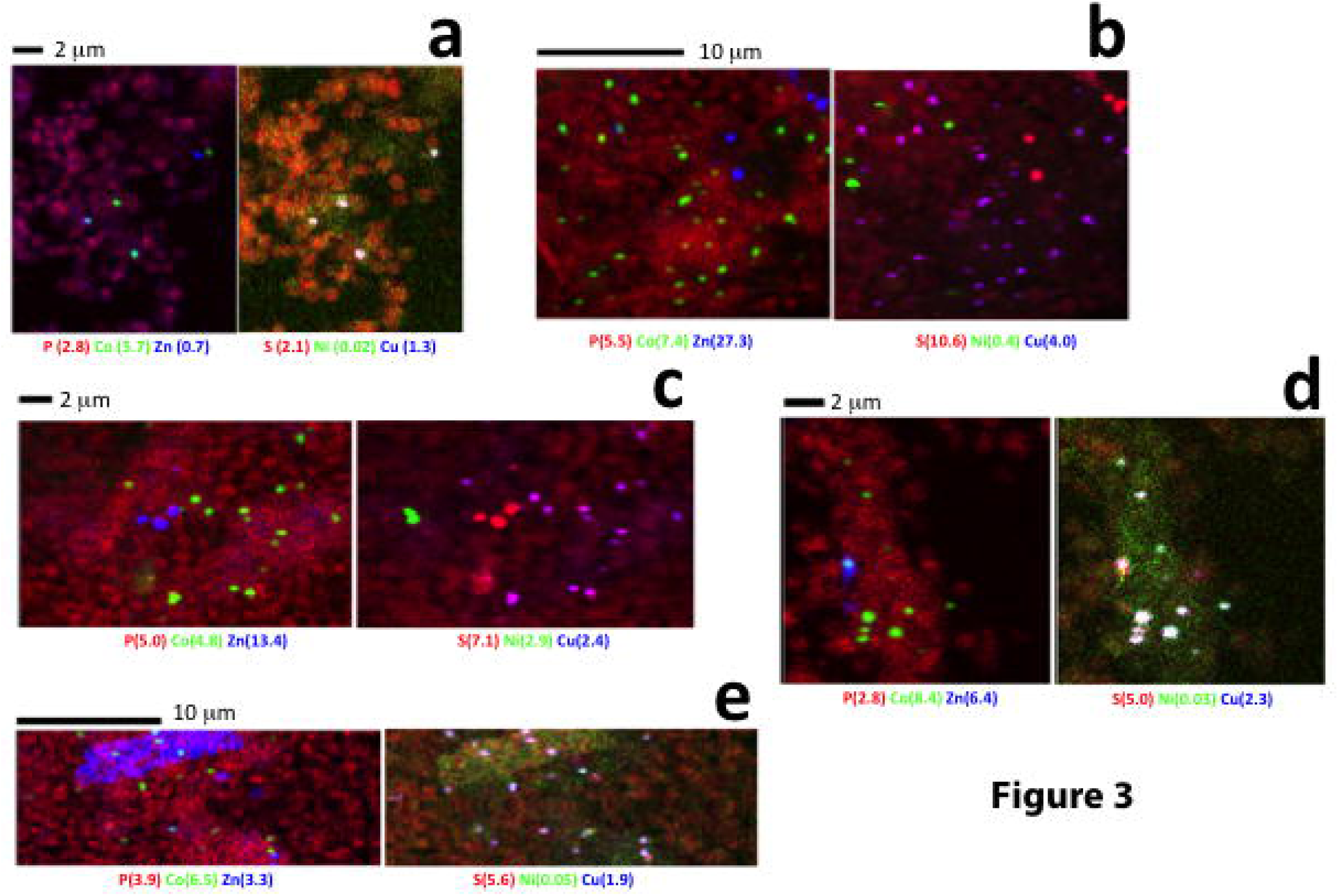
SXRF co-localization of P (red), Co (green), and Zn (blue) in left panels, and S (red), Ni (green), and Cu (blue) in right panels for five imaged fields of 5 μm thin sections of 77% *Methanosarcina acetivorans* and 23% *Desulfococcus multivorans* cocultures. Values in parentheses are maxima in g cm-2 for each element.

## Discussion

In this study, SXRF imaging and quantification of trace metals in cellular and abiotic phases was performed at the single-cell scale. Our observation that Zn and Fe were the two most abundant cellular trace metals in monocultures is consistent with previous studies of diverse prokaryotes (Barton et al. 2007; Cvetkovic et al. 2010; Outten and O’Halloran 2001; Rouf 1964), including diverse mesophilic and hyperthermophilic methanogens grown on a range of substrates, for which, generally: Fe > Zn > Ni > Co > Cu (Cameron et al. 2012; Scherer et al. 1983). To our knowledge, there are no previous reports of the trace metal content of sulfate-reducing bacteria, but the abundance of Fe and Zn-containing proteins encoded by their genomes (Barton and Fauque 2009; Barton et al. 2007; Fauque and Barton 2012) is consistent with the cellular enrichment we observed in these trace metals.

Both normalizations for SXRF data (per cell and per cellular volume) showed that the methanogenic archaeon contained more P, S, Co, Ni and Cu than the sulfate-reducing bacterium. The higher cellular Co content of *M. acetivorans* vs. *D. multivorans* is likely due to due to numerous methyltransferases involved in methylotrophic methanogenesis (Zhang and Gladyshev 2010; Zhang et al. 2009) that contain cobalt as a metal center in their corrinoid (vitamin B12) cofactor, in addition to the corrinoid-containing Fe-S methyltransferase protein in the Wood Ljungdahl pathway in both species (Ekstrom and Morel 2008; Fig.4). Similarly, the higher Ni content of *M. acetivorans* vs. *D. multivorans* is likely due to the presence of Ni-containing cofactor F430 in methyl coenzyme M reductase, the final enzyme in the methanogenesis pathway. Cofactor F430 is found only in methane-metabolizing archaea, in which it comprises 50-80% of total cellular Ni (Diekert et al. 1981; Mayr et al. 2008). Additional Ni requirements in both *M acetivorans* and *D. multivorans* are used for Ni-Fe hydrogenases and carbon monoxide dehydrogenase in the Wood-Ljungdahl pathway (Fig.4).

**Figure 4.**
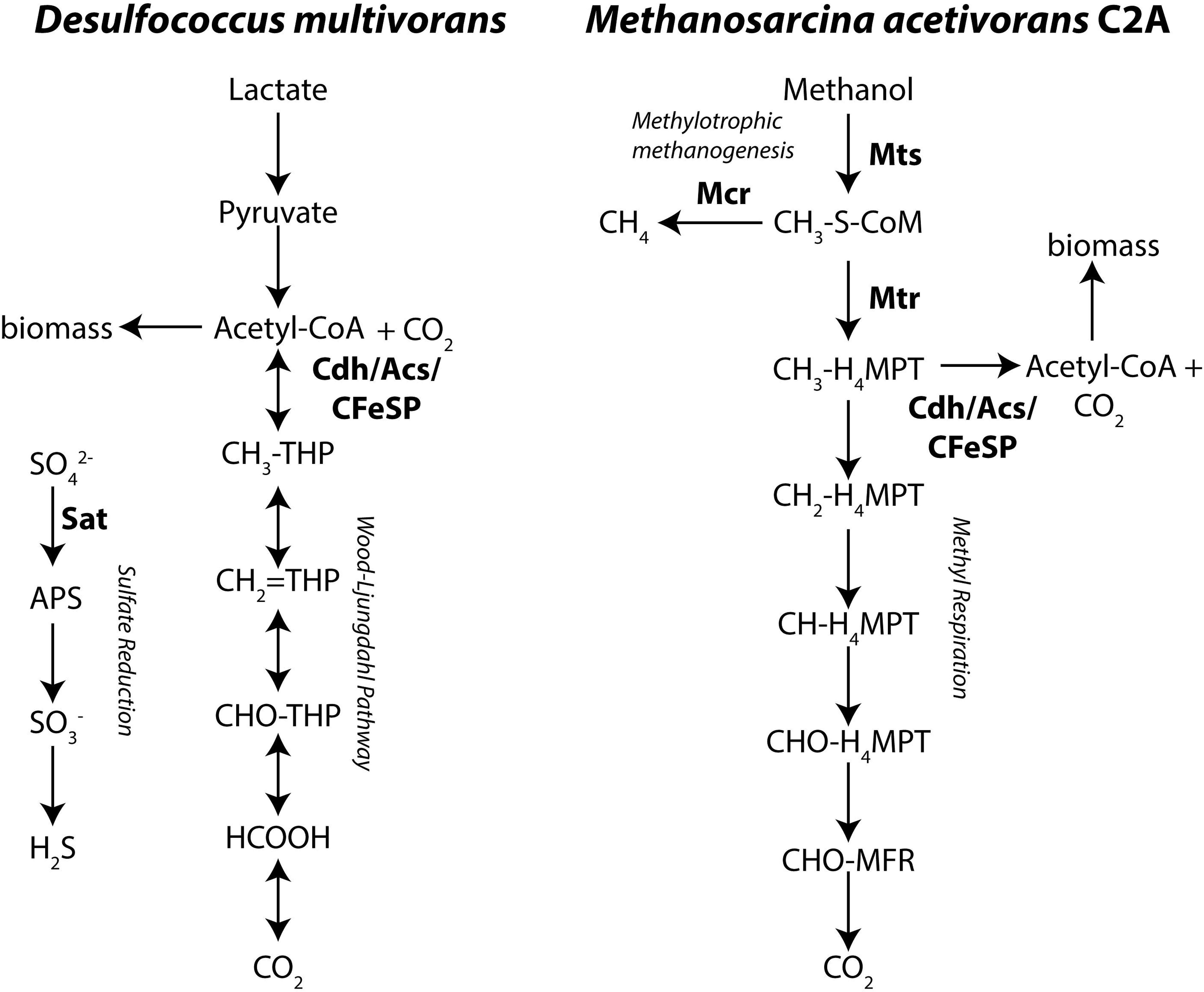
Schematic of metalloenzyme-containing metabolic pathways in the complete carbon-oxidizing sulfate-reducing bacterium *Desulfococcus mutitvorans* and the methylotrophic methanogenic archaeon *Methanosarcina acetivorans* as confirmed by genomic analyses. Nickel (Acs, Cdh, Mcr) and cobalt (CFeSP, Mts, Mtr, and Sat) containing enzymes are labeled in bold. Enzyme abbreviations: Acs/CFeSP: acetyl-CoA synthase/corrinoid-FeS protein; Cdh: carbon monoxide dehydrogenase; Mts: methanol:coenzyme M methyltransferase; Mcr: methyl coenzyme M reductase; Mtr: methyl-tetrahydromethanopterin:coenzyme M methyltransferase; Sat: ATP sulfurylase.

Metabolic Cu requirements for methanogenesis are not well known, although high accumulations have also been reported for other methanogens (Scherer et al. 1983). However, it should be noted that our early trials analyzing S-rich cells on Au grids revealed artifacts resulting from interactions of S and Cu underlying the grid’s surface Au coating (data not shown); use of SiN grids in this study appeared to eliminate such Cu artifacts, but potential reactions between trace Cu in SiN grids and abundant S in the archaeal cells cannot be completely discounted.

Faster growth rates of methylotrophic methanogens than sulfate-reducing bacteria at moderate temperatures have been reported in previous studies (Dawson et al. 2015; Weijma and Stams 2001), and likely account for *M. acetivorans* outcompeting *D. multivorans* in our cocultures. We consider it unlikely that differences in cellular trace metal contents in monocultures were a result of harvesting *D. multivorans* earlier in their stationary phase than *M.acetivorans* (Fig. S1) because cellular metal reserves generally decline or remain constant in stationary phase (Bellenger et al. 2011). Our SXRF measurements of cocultures are more difficult to interpret due to apparent swelling of aldehyde-fixed cocultured cells (∼1 μm radius) to ∼2x the size of monocultures (0.5-0.6 μm radius). When normalized per cell, fixed cocultures showed significantly higher P, Co and Ni than unfixed monocultures, but the apparent swelling of cocultured cells erased this trend when normalized to cellular volume.

When grown at millimolar metal concentrations, sulfate-reducing bacteria efficiently remove metals from solution (Krumholz et al. 2003) and precipitate covellite (CuS; Gramp et al. 2006; Karnachuk et al. 2008), sphalerite/wurtzite (ZnS/(Zn,Fe)S; Gramp et al. 2007; Xu et al. and pentlandite (Co_9_S_8_) (Sitte et al. 2013). Based on its ∼1:1 Zn:S ratio, the semi-circular nanoparticulate zinc sulfide phase(s) observed in thin sections imaged by SXRF in this study were likely sphalerite spheroids, also found in sulfate-reducing bacteria biofilms due to aggregation of ZnS nanocrystals (0.1-10 μm) and extracellular proteins (Moreau et al. 2004; Moreau et al. 2007). The abiotic phase with the approximate stoichiometry (CoCu)S_2_ may be mineralogically distinct from those in previous studies.

## Conclusions and Challenges

This study used two independent methods for assessing trace metal inventories in anaerobic microbial cultures. We found that SXRF is a promising method for imaging and quantifying first-row transition metals in anaerobic microbial cultures at single-cell resolution. This method’s single-cell resolution enables more precise measurements of cellular metal content than ICP-MS analysis of bulk cells, which can include metals bound to extracellular aggregations such as cation-binding exopolymeric substances produced by sulfate-reducing bacteria (Beech and Cheung 1995; Beech et al. 1999; Braissant et al. 2007). We did not observe evidence of metal contamination from aldehyde fixation in SXRF data, likely because we pre-cleaned fixatives with metal-chelating resin prior to use, as previously described by Twining et al. (2003).

Challenges remain with accurate elemental quantification of microbial cocultures preserved in a manner that would also allow assignment of identity for similar cell types. It was not possible to distinguish methanogenic archaea from sulfate-reducing bacteria in coculture on the basis of cell morphology or elemental content, and attempts to image cells with fluorescent oligonucleotide probes after SXRF analysis were unsuccessful due to x-ray radiation damage. We recommend method development for simultaneous taxonomic identification and elemental imaging (e.g. gold-FISH (Schmidt et al. 2012)) for samples containing multiple microbial species as a high priority for future work

## Author Contributions

J.B.G., V.J.O., S.C., and K.S.D. conceived and designed the experiments; K.S.D. performed the microbial culturing, S.C., J.B.G., and S.V. performed the SXRF analyses; B.S.T. performed the ICP-MS analysis, J.B.G., S.C., D.R.H., S.V., E.D.I., and B.S.T. analyzed the data; and J.B.G. wrote the manuscript with input from all authors. All authors have given approval to the final version of the manuscript.

## Acknowledgements

We thank Shawn McGlynn for assistance with sample preparation, and Keith Brister, Junjing Deng, Barry Lai and Rachel Mak at LS-CAT for assistance with Bionanoprobe analysis. We thank Larry Barton and Joel Kostka for helpful discussions, and Marcus Bray, Amanda Cavazos, Grayson Chadwick, Chloe Stanton, and Nadia Szeinbaum for feedback on previous manuscript drafts.

---

**Figure S1.** Growth curves based on OD600 for the three cultures described in this study: *Methanosarcina acetivorans* (white), *Desulfococcus multivorans* (light grey), and 77% *Methanosarcina acetivorans* and 23% *Desulfococcus multivorans* cocultures (dark grey).

